# Inferring Population Structure and Admixture Proportions in Low Depth NGS Data

**DOI:** 10.1101/302463

**Authors:** Jonas Meisner, Anders Albrechtsen

## Abstract

We here present two methods for inferring population structure and admixture proportions in low depth next generation sequencing data. Inference of population structure is essential in both population genetics and association studies and is often performed using principal component analysis or clustering-based approaches. Next-generation sequencing methods provide large amounts of genetic data but are associated with statistical uncertainty for especially low depth sequencing data. Models can account for this uncertainty by working directly on genotype likelihoods of the unobserved genotypes. We propose a method for inferring population structure through principal component analysis in an iterative approach of estimating individual allele frequencies, where we demonstrate improved accuracy in samples with low and variable sequencing depth for both simulated and real datasets. We also use the estimated individual allele frequencies in a fast non-negative matrix factorization method to estimate admixture proportions. Both methods have been implemented in the PCAngsd framework available at http://www.popgen.dk/software/.

---

POPULATION genetic studies often consist of individuals of diverse ancestries, and inference of population structure therefore plays an important role in population genetics and association studies. Population stratification can act as a confounding factor in association studies as it can lead to spurious associations (Marchini *et al.* 2004). Principal component analysis (PCA) was first introduced to genetic data in Menozzi *et al.* (1978) to produce synthetic maps in an exploratory analysis of genetic variation. PCA is now a common tool in population genetic studies, where its dimension reduction properties can be used to visualize population structure by summarizing the genetic variation through principal components (Novembre and Stephens 2008), correct for population stratification in association studies, investigate demographic history (Patterson *et al.* 2006; Fumagalli *et al.* 2013; Price *et al.* 2006) as well as perform genome selection scans (Galinsky *et al.* 2016; Hao *et al.* 2015; Luu *et al.* 2017). PCA is an appealing approach to infer population structure as the aim is not to classify the individuals into discrete populations, however instead describe continuous axes of genetic variation such that heterogeneous populations and admixed individuals can be better represented (Patterson *et al.* 2006). Another successful approach in modeling complex population structure is to estimate admixture proportions based on clustering-based methods (Pritchard *et al.* 2000; Tang *et al.* 2005; Alexander *et al.* 2009; Skotte *et al.* 2013), such as the popular software ADMIXTURE, which have also been used for correction of population stratification in association studies (Price *et al.* 2010).

Next-generation sequencing (NGS) methods (Metzker 2010) produce a large amount of DNA sequencing data at low cost and are commonly used in population genetic studies (Nielsen *et al.* 2012). But NGS methods are associated with high error rates usually caused by several factors such as sampling, alignment and sequencing errors. Many NGS studies are based on medium (*<*15X) and low (*<*5X) depth data due to the demand for large sample sizes as seen in large-scale sequencing studies, e.g. 1000 Genomes Project Consortium (Consortium *et al.* 2010, 2012). However, the use of medium and especially low depth sequencing data introduces challenges rooted in the statistical uncertainty induced when calling genotypes and variants in these scenarios (Nielsen *et al.* 2012). The statistical uncertainty increases for low depth samples due to the increased difficulty of distinguishing a variable site from a sequencing error with the information provided. Problems can arise due to chromosomes being sampled with replacement in the sequencing process, and both alleles may not have been sampled for a heterozygous individual in low depth scenarios. Homozygous genotypes may also be wrongly inferred as heterozygous due to sequencing errors. Thus, genotype calling will associate individuals with a statistical uncertainty which should be taken into account (Nielsen *et al.* 2011, 2012).

To overcome these problems related to NGS data and genotype calling, probabilistic methods have been developed to take use of genotype likelihoods in combination with external information for various population genetic parameters (Kim *et al.* 2011; Nielsen *et al.* 2012; Korneliussen *et al.* 2014; Skotte *et al.* 2013; Fumagalli *et al.* 2013; Vieira *et al.* 2013; Kousathanas *et al.* 2017), such that posterior genotype probabilities can be used to model the related uncertainty. Genotype likelihoods can be estimated to incorporate errors of the sequencing process such as the base quality scores as well as the allele sampling (McKenna *et al.* 2010). These posterior genotype probabilities have also been used to call genotypes with a higher accuracy than previous methods for low depth NGS data (Nielsen *et al.* 2011, 2012).

We present two new methods for low depth NGS data using genotype likelihoods to model complex population structure that connect the results of PCA with the admixture proportions of clustering-based approaches. A method has been developed to perform PCA in an iterative approach of estimating individual allele frequencies to compute a covariance matrix, while another method uses the estimated individual allele frequencies in an accelerated non-negative matrix factorization (NMF) algorithm to estimate admixture proportions. The performances of the two methods are assessed on both simulated and real datasets in regards to existing methods for both low depth NGS and genotype data. The methods have been implemented in a framework called PCAngsd (Principal Component Analysis of NextGeneration Sequencing Data).

## Materials and Methods

We will analyze NGS data of *n* diploid individuals across *m* variable sites. These sites will either be known or called single nucleotide polymorphisms (SNPs), which are assumed to be diallelic such that the major and minor allele of each SNP have been inferred. This can either be done from sequencing reads (Kim *et al.* 2011) or from genotype likelihoods (Korneliussen *et al.* 2014) and only three different genotypes will be possible. Thus, we assume that a genotype *G* can be seen as a Binomial random variable with realizations 0, 1 and 2 that represent the number of copies of the minor allele in a site for a given individual in the absence of population structure. The expectation and variance of *G* can therefore be defined as 𝔼 [*G*] = 2*p* and Var[*G*] = 2*p*(1 − *p*) with *p* representing the allele frequency of a population, which we also refer to as population allele frequency.

However, genotypes are not observed in NGS data and we will instead work on genotype likelihoods that also include information of the sequencing process. The genotype likelihoods are the probability of the observed sequencing data *X* given the three different possible genotypes, *P(X*|*G* = *g*), for *g* = 0, 1, 2. One method to compute genotype likelihoods from sequencing reads is described in the supplementary material based on the simple GATK model (McKenna *et al.* 2010).

External information can be incorporated to define posterior genotype probabilities using Bayes′ theorem in combination with genotype likelihoods (Nielsen *et al.* 2011). The population allele frequency is often used as information in the estimation of prior genotype probability *P*(*G* _*is*_ | *p*_*s*_), for an individual *i* in site *s* (Kim *et al.* 2011; Nielsen *et al.* 2012; Fumagalli *et al.* 2013; Vieira *et al.* 2013). Assuming the population is in Hardy-Weinberg Equilibrium (HWE) for a site *s*, the prior genotype probability is then given as *P*(*G*_*is*_ = 0| *p*_*s*_) = (1 − *p*_*s*_)^2^, *P*(*G*_*is*_ = 1| *p*_*is*_) = 2*p*_*s*_ (1*p*_*s*_) and 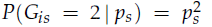 for the three different possible genotypes. As defined in Kim *et al.* (2011), using the estimated population allele frequency *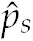,* the posterior genotype probability is computed as follows for individual *i* in site *s*:

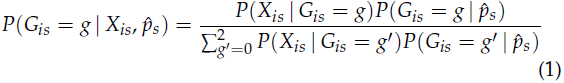

### PCA

The standard way of performing PCA in population genetics and using it to infer population structure is based on the method defined in Patterson *et al.* (2006). For a genotype matrix **G** of *n* individuals and *m* variable sites, the *n* × *n* covariance matrix **C**, also known as the genetic relationship matrix (GRM), is computed as follows for two individuals *i* and *j*:

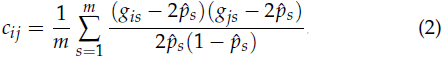

Here *g*_*is*_ is the observed genotype for individual *i* in site *s* to distinguish it from *G* defined above for unobserved genotypes, and 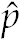 is the estimated population allele frequency. The principal components are then inferred by performing an eigendecomposition of the covariance matrix, such that **C** = **VΣV^*T*^** with **V** being the matrix of eigenvectors and **Σ** the diagonal matrix of the corresponding eigenvalues. Principal components and eigenvectors will be used interchangeably throughout this study. The top principal components capture most of the population structure as they represent axes of genetic variation in the dataset (Patterson *et al.* 2006).

This method has been extended to NGS data in Fumagalli *et al.* (2013), as well as in Skotte *et al.* (2012), using the probabilistic framework described in equation 1, by summing over the genotypes of each individual weighted by the joint posterior genotype probabilities under the assumption of HWE in the whole sample. The method has been implemented in the ngsTools framework (Fumagalli *et al.* 2014). The covariance matrix is estimated as follows for NGS data using only known variable sites for two individuals *i* and *j*:

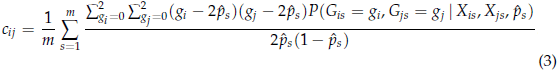

ngsTools splits up the joint posterior probability, 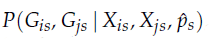, into 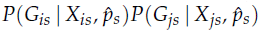 for *i* ≠ *j* by assuming conditional independence between individuals given the estimated population allele frequencies. The non-diagonal entries in the covariance matrix are now directly estimated from the posterior expectations of the genotype instead of the observed genotypes as described in equation 2. The original method weighs each site by its probability of being a variable site such that SNP calling is not needed prior to the covariance matrix estimation. This is not taking into account in this study as we are using called variable sites to infer population structure. The population allele frequencies are estimated from the genotype likelihoods using an expectation maximization (EM) algorithm (Kim *et al.* 2011) as described in the supplementary material.

The problem with this approach is that the assumption of conditional independence between individuals given the population allele frequency is only valid when there is no population structure. Here we propose a novel approach of estimating the covariance matrix using iteratively estimated individual allele frequencies to update the prior information of the posterior genotype probability. Thereby we condition on the individual allele frequencies as in the clustering-based approaches.

### Individual allele frequencies

A model for estimating individual allele frequencies based on population structure was introduced in STRUCTURE (Pritchard *et al.* 2000) as later described in equation 13. Hao et al. proposed a different model for estimating individual allele frequencies Π by using the information in the principal components instead of having an assumption of *K* ancestral populations (Hao *et al.* 2015). The model is defined as the matrix product,

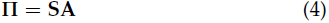

where **S** represents the population structure such that **A** represents the mapping of the population structure **S** to the allele frequencies. Hao et al. estimated the individual allele frequencies through a singular value decomposition (SVD) method, where genotypes are reconstructed using only the top *D* principal components such that they will be modeled by population structure. A similar approach has been proposed by Conomos et al. (Conomos *et al.* 2016) where the inferred principal components are used to estimate individual allele frequencies in a simple linear regression model. However, due to working on NGS data and not knowing the genotypes, we are extending the method of Hao et al. to NGS data by using posterior expectations of the genotypes, referred to as genotype dosages, instead of genotypes. Thus we will be using,

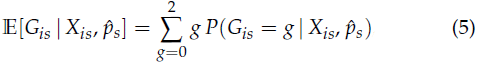

for individual *i* in site *s*

The individual allele frequencies are then estimated by performing a SVD on the centered genotype dosages and reconstructing them using only the top *D* principal components. 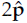 is then added to the reconstruction and scaled by 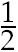 based on a Binomial distribution assumption of *G*_*s*_, for *i* = 1,…, *n* and *s* = 1,…, *m*, to produce the individual allele frequencies. Since SVD is a real valued method, we will have to truncate the estimated individual allele frequencies in order to constrain them in the range [0, 1]. However, Hao et al. showed that the resulting estimates were still very accurate for common variants considering this limitation.

For ease of notation, let **E** be the *n* × *m* matrix of genotype dosages, 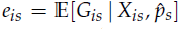 for *i* = 1,…,*n* and *s* = 1,…, *m*. The following steps for estimating the individual allele frequencies are adopted from the SVD method (Hao *et al.* 2015) to work on NGS data:

#### Algorithm 1: SVD method for estimating individual allele frequencies

1. The centered genotype dosages are constructed as 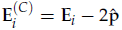 for *i* = 1,…, *n*.

2. Perform SVD on the centered genotype dosages, **E**^*(C)*^ = **WΔU**^*T*^, where **W** will represent population structure similarly to **V**.

3. Define 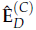 to be the prediction of the centered genotype dosages using only the top *D* principal components, 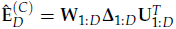

4. Estimate 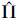 by adding 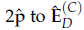 row-wise and scaling by 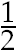, based on 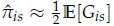.

For matrix notations define 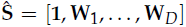 and 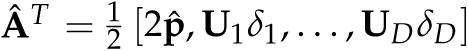 all representing column vectors, such that equation 4 can be approximated as. 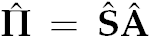 Finally, 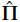 is truncated to constrain allele frequency estimates in a range based on a small value *γ* (1.0 × 10^−4^), such that 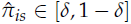 for *i* = 1,…, *n* and *s* = 1,…, *m*.

We now incorporate the individual allele frequencies into the estimation of posterior genotype probabilities. The estimated individual allele frequencies are used as updated prior information instead of the population allele frequencies. The individual allele frequencies will then be able to model missing data with better estimates of the genotypes given the inferred population structure of the individuals. Thus, the posterior genotype probabilities are estimated as follows: for individual *i* in site *s*

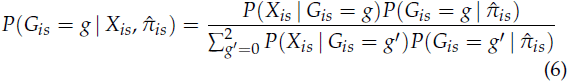

Each individual are now seen as a single population with allele frequency 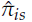 where as the prior genotype probability are estimated assuming HWE, such that 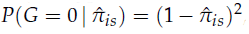 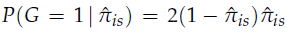 and 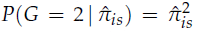. An updated definition of the posterior expectations of the genotypes are then given as:

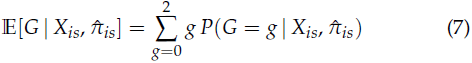

This procedure of updating the prior information can be iterated to estimate new individual allele frequencies on the basis of updated population structure. Therefore, we propose the following algorithm for an iterative procedure of estimating the individual allele frequencies.

#### Algorithm 2: Iterative estimation of individual allele frequencies

1. Estimate population allele frequencies 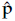 from genotype likelihoods (See supplementary materials).

2. Estimate posterior genotype probabilities and genotype dosages **E** based on genotype likelihoods and 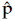

3. Estimate 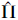 using the SVD based method on **E** as described in Algorithm 1.

4. Estimate posterior genotype probabilities and genotype dosages **E** using updated prior information, 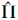.

5. Repeat step 3 and 4 until individual allele frequencies have converged.

Convergence of our iterative method is defined as when the root-mean-square deviation (RMSD) of the inferred population structure in the SVD **W** are smaller than a value *δ* (1.0 × 10^−5^) between two successive iterations. The RMSD of iteration *t* + 1 for *D* principal components is given as,

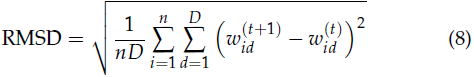

### Covariance matrix

We now use the final set of individual allele frequencies to estimate an updated covariance matrix in a similar model as in equation 3, but incorporating the individual allele frequencies into the joint posterior probability. The entries of the covariance matrix **C** are now defined as follow for individuals *i* and *j*:

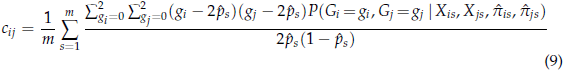

For *i ≠ j*, the joint posterior probability can be computed as 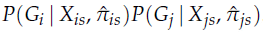, since the individuals are conditionally independent given the individual allele frequencies in contrary to the assumption made in the model of Fumagalli et al. using population allele frequencies. The above equation can be expressed in terms of the genotype dosages for ease of notation and computation:

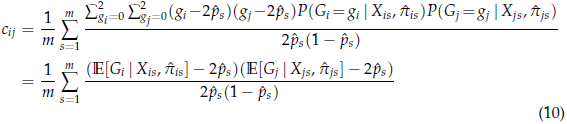

However for *i* = *j* (diagonal of the covariance matrix), the joint posterior probability is simplified to 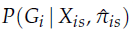 such that the estimation of the diagonal covariance entries is given as:

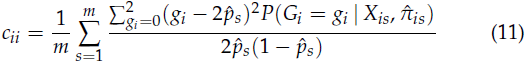

An eigendecomposition of the updated estimated covariance matrix is then performed to obtain the principal components as described earlier, **C** = **VΣV**^*T*^. Note that **V** and **W** from algorithm 1 are not the same even though they both represent population structure through axes of genetic variation in the dataset. This is due to a different scaling and the joint posterior probability of equation 11 is not taken into account in **W** for i = j.

### Number of principal components

It can be hard to determine the optimal number of principal components that represent population structure. In our method, we are using Velicier’s minimum average partial (MAP) test as proposed by Shriner (Shriner 2011) to automatically detect the number of top principal components *D* used for estimating the individual allele frequencies. Shriner showed that the test based on a Tracy-Widom distribution (Patterson *et al.* 2006) systematically overestimates the number of significant principal components and even performs worse for datasets including admixed individuals. However, in order to be able to perform the MAP test and detect the optimal *D*, an initial covariance matrix is estimated based on the model in equation 3.

The MAP test is performed on the estimated initial covariance matrix **C** for NGS data as an approximation of the Pearsson correlation matrix used by Shriner. Using the notation of Shriner, 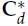 is defined as the matrix of partial correlations after having partialed out the first *d* principal components. Velicer (Velicer 1976) proposed the summary statistic 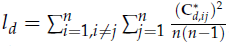 where 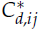 represents the entry in 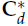 for individuals *i* and *j* Thus, the test statistic*l* _*d*_ represents the average squared correlation after partialing out the top *d* principal components. The number of top principal components that represent population structure is then chosen as argmin_*d*_ *l*_*d*_, for *d* = 0,… *m –* 1. We have used the same implementation of the MAP test as Shriner.

The MAP test is performed on the estimated initial covari ance matrix can be avoided by having prior knowledge of an optimal *D* for the dataset being analyzed and manually selecting *D*

### Genotype calling

As previously shown in Nielsen *et al.* (2012); Fumagalli *et al.* (2013), genotypes can be called from posterior genotype probabilities to achieve higher accuracy in low depth NGS scenarios. We can adapt this concept to our posterior genotype probabilities based on individual allele frequencies, such that genotypes can be called at a higher accuracy in structured populations from low depth NGS data. The genotype for individual *i* in site *s* is called as follows:

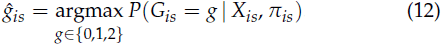

### Admixture proportions

Based on the likelihood model defined in STRUCTURE (Pritchard *et al.* 2000), individual allele frequencies **Π** can be estimated using admixture proportions **Q** and population-specific allele frequencies **F** (Alexander *et al.* 2009), such that:

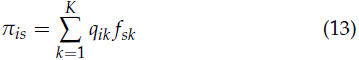

for an individual *i* in a variable site *s*. This is based on an assumption of *K* ancestral populations where 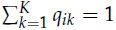 and 0 *≤ q, f ≤* 1 ∀ *q, f ∈* (**Q**, **F**). Here **Q** and **F** must be inferred in order to estimate the individual allele frequencies, where as *K* is assumed to be known. One probabilistic approach for inferring population structure through admixture proportions for low depth NGS data has been implemented in the NGSadmix software (Skotte *et al.* 2013). Here both parameters, **Q** and **F**, are jointly estimated in an EM algorithm using genotype likelihoods

In our case, we have already estimated the individual allele frequencies based on our iterative procedure using PCA described above. *K* can be chosen as the number of principal components *D* + 1, since it would explain the number of distinct ancestral population from which the individual allele frequencies have been estimated from. There is however not always a direct interpretation between principal components and admixture proportions (Alexander *et al.* 2009). Therefore, we propose an approach based on non-negative matrix factorization (NMF) to infer **Q** and **F** using only our estimated individual allele frequencies as information for low depth NGS data. NMF has previously been applied directly on genotype data to infer population structure and admixture proportions by Frichot et al. (Frichot *et al.* 2014), where their method showed comparable accuracy and faster run-time in comparison to ADMIXTURE

NMF is a dimension reduction and factor analysis method for finding a low-rank approximation of a matrix, which is similar to PCA, but NMF is constrained to find non-negative low dimensional matrices. For an non-negative matrix 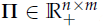 the goal of NMF is to find an approximation of **Π** based on two non-negative factor matrices 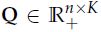 and 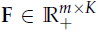 such that

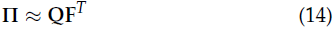

**Q** will consist of columns of non-negative basis vectors such that linear combinations of these approximates **Π** through **F**. Thus based on the non-negative nature of our parameters, we can apply the ideas of NMF to infer admixture proportions **Q** and population-specific allele frequencies **F** from our individual allele frequencies. We use a combination of recent research in NMF to minimize the following least squares problem with a sparseness constraint on **Q**:

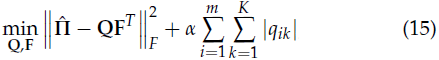

for **Q** ≥ 0, **F** ≥ 0 and *α* ≥ 0. Here ‖. ‖ _*F*_ is the Frobenius norm of a matrix and *α* is the regularization parameter controlling the sparseness enforced as also introduced in Frichot *et al.* (2014).

Lee and Seung (Lee and Seung 1999, 2001) proposed an multiplicative update (MU) algorithm to solve the standard NMF problem without the sparseness constraint included above. Their update rules can be seen as conservative steps in a gradient descent optimization problem for updating **F** and **Q**, which ensure that the non-negative constraint holds for each update. Hoyer (Hoyer 2002) extended the MU to incorporate the sparseness constraint described in equation 15 for **Q**. For *α >* 0, the regularization parameter is used to reduce noise, especially induced by the uncertainty of low depth NGS data, in the estimated admixture proportions by enforcing sparseness in the solution. An iteration of using the MU rules are then described as follows:

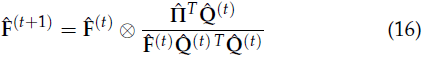

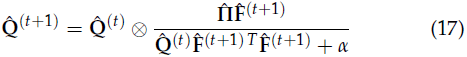

where ⊗ represents element-wise multiplication and the division operator is elementwise as well.

However, MU has been shown to have a slow convergence rate, especially for dense matrices, and our approach is therefore to accelerate MU by combining two different techniques. We propose an algorithm of combining the acceleration scheme described by Gillis and Glineur (Gillis and Glineur 2012) with the asymmetric stochastic gradient descent algorithm (ASG-MU) of Serizel et al. (Serizel et al. 2016) for updating **F** and **Q** in a fast approach. The acceleration scheme of Gillis and Glineur (2011) updates each matrix **F** and **Q** a fixed number of times at a lower computational cost without losing the convergence properties of MU.We simply incorporate this acceleration scheme inside ASGMU that works by randomly assigning the columns **Π** of into a set of B mini-batches, which are then updated sequentially in a permuted order to improve the convergence rate and performance of MU (Serizel et al. 2016). After each update, we truncate the entries of both **F** and **Q** to be in range [0, 1] and normalize the rows of **Q** to sum to one. The concept of combining an acceleration scheme with a stochastic gradient descent approach for MU has also been explored in Kasai (2017).

The algorithm is iterated until the admixture proportions has converged. Convergence is defined as when the RMSD of estimated admixture proportions of two successive iterations are smaller than a value *φ* (1.0 × 10^−4^). The RMSD of iteration *t* + 1 is given as,

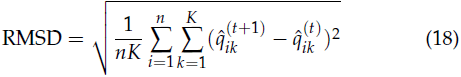

The *α* parameter enforcing sparseness in the estimated solution of **Q** is arbitrarily specified. However the use of the likelihood measure in the NGSdamix (Skotte *et al.* 2013) model can be used to determine the *α* parameter fitting the dataset. The likelihood measure is defined as:

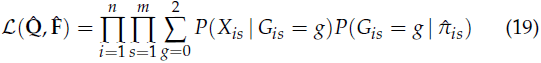

Where 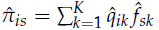. Based on the fast estimation of admixture proportions using our NMF algorithm, a set of *α* values can be tested and measured sequentially using the likelihood measure. This can be performed without sacrificing significant run-time compared to NGSadmix due to already having estimated the individual allele frequencies for a particular *K*.

### Implementation

Both presented methods have been implemented in a Python framework named PCAngsd (Principal Component Analysis of Next-Generation Sequencing Data). The framework is freely available at http://www.popgen.dk/software/.

The memory requirements of PCAngsd is 𝒪(*mn*) as the entire matrix of genotype likelihoods needs to be stored in memory for both methods. The most computational expensive step is the estimation of individual allele frequencies and covariance matrix (𝒪 (*m*^2^*n*)). However, a fast SVD method for only computing the top *D* eigenvectors, implemented in the Scipy library (Jones *et al.* 2014) using ARPACK (Lehoucq *et al.* 1998) as an eigensolver, has been used to speed up the iterative estimations of the individual allele frequencies. PCAngsd is multithreaded as well to take advantage of several cores and the backbone of the framework is based on Numpy data structures (Walt *et al.* 2011) using the Numba library (Lam *et al.* 2015) to speed up bottlenecks with just-in-time (JIT) compilation.

### Simple simulation of genotypes and sequencing data

Low depth NGS data has been simulated as genotype likelihoods to test the capabilities of our two presented methods. Allele frequencies of the reference panel of the Human Genome Diversity Project (HGDP) (Cann *et al.* 2002) have been used to generate a total of 380 individuals from three distinct populations (French, Han Chinese, Yoruba) including admixed individuals in approximately 0.4 million SNPs across all autosomes. As the allele frequencies are known for each population, the genotypes of each individual can be sampled from a Binomial distribution for each diallelic SNP, using the population-specific allele frequency or an admixed allele frequency as parameter. No LD has been simulated. The genotypes are therefore known and are used in the evaluation of our methods in our low depth scenarios. The number of reads in each SNP are sampled from a Poisson distribution with a mean parameter resembling the average sequencing depth of the individual, and the genotype is used to sample the number of derived alleles from a Binomial distribution using the sampled depth as parameter. The average sequencing depth of each individual is sampled uniformly random from a range of [0.5, 5]. Sequencing errors are incorporated by sampling each read with a probability *ϵ* = 0.01 of being an error. The genotype likelihoods are then finally generated from the probability mass function of a Binomial distribution using the sampled parameters and *e*. This approach of genotype likelihood simulation has previously been used in Kim *et al.* (2011); Vieira *et al.* (2013); Skotte *et al.* (2013).

A complex admixture scenario has been constructed to test the capabilities of our methods. 100 individuals have been sampled directly from each of the population-specific allele frequencies (non-admixed), while 50 individuals have been sampled to have equal ancestry from each of the three distinct popula tions (threeway admixture). At last, 30 individuals have been sampled from a gradient of ancestry between all pairs of the ancestral populations (two-way admixture).

### 1000 Genomes low depth sequencing data

We also analyze human low coverage NGS data of 193 individuals from the 1000 Genomes Project Consortium (Consortium *et al.* 2010, 2012). The individuals are from four different populations consisting of 41 from CEU (Utah residents with Northern and Western European ancestry), 40 from CHB (Han Chinese in Beijing), 48 from YRI (Yoruba in Ibadan) and 64 individuals from MXL (Mexican ancestry in Los Angeles) representing an admixed scenario of European and Native American ancestry. The individuals from the low coverage datasets have a varying sequencing depth from 1.5 − 12.5 𝒳 after site filtering. An advantage of using the low coverage data of the 1000 Genomes Project data is that overlapping reliable genotypes are available which can be used for validation purposes.

SNP calling and estimation of genotype likelihoods of the 1000 Genomes dataset has been performed in ANGSD (Kor neliussen *et al.* 2014) using simple read quality filters. A significance threshold of 1.0 × 10^−6^ has been used for SNP calling alongside a MAF threshold of 0.05 to remove rare variants. A total number of 8 million variable sites across all autosomes have been used in the analyses. The full ANGSD command used to generate the genotype likelihoods is provided in the supplementary material.

### Waterbuck low depth sequencing data

Lastly, an animal dataset (non-model organism) has also been included in our study. A reduced low depth NGS dataset of the waterbuck (*Kobus ellipsiprymnus*) originating from Pedersen et al. (unpublished) has been analyzed. The dataset consists of 73 samples that have been sampled at 5 different sites in Africa with a varying sequencing depth from 2.2 − 4.7 𝒳 aligned to 730 scaffolds. The dataset has been reduced to only include sampling sites with more than 10 samples such that the inferred axes of genetic variation will reflect true population structure. As performed for the 1000 Genomes dataset, genotype likelihoods has been estimated in ANGSD with the same SNP and MAF filters. A total number of 9.4 million SNPs across the autosomes of the waterbuck is analyzed in this study.

## Results

For the simulated and 1000 Genomes datasets, results estimated in PCAngsd on low depth NGS data are evaluated against the results estimated from genotype data. The model in equation 2 is used to perform PCA, while ADMIXTURE is used to estimate admixture proportions on the “true” genotype datasets. The performance of PCAngsd is also compared to existing genotype likelihood methods with the ngsTools model (equation 3) for performing PCA, and NGSadmix (equation 19) for estimating admixture proportions. In all the following cases of admixture plots estimated by PCAngsd, we have used *B* = 5 and *α* has been chosen as the one maximizing the likelihood measure described above (equation 19), also shown in Figure S5.

RMSD is used to evaluate the performances of both NGS methods for estimating admixture proportions in terms of accuracy:

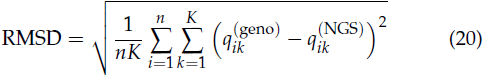

Where 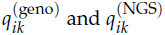 and represents the estimated admixture proportion for individual *i* in ancestral population *k* from known genotypes and NGS data, respectively. The accuracy of the inferred PCA plots of both NGS methods are also compared to the PCA plots of the known genotypes for the simulated and 1000 Genomes datasets using RMSD. However, a Procrustes analysis (Wang *et al.* 2010; Fumagalli *et al.* 2013) must be performed prior to the comparison as the direction of the principal components can differ based on the eigendecomposition of the covariance matrices.

All tests in this study have been performed serverside using 32 threads (Intel® Xeon® CPU E5-2690) for both PCAngsd and NGSadmix.

### Simulation

The results of performing PCA on the simulated dataset based on frequencies from 3 human populations are displayed in Figure 1, where we simulated unadmixed, two-way admixed and three way admixed individuals. The MAP test reported 2 significant principal components which was also expected for individuals simulated from three distinct populations. The inferred principal components clearly shows the importance of taking individual allele frequencies into account in the probabilistic framework. Here PCAngsd is able to infer the population structure of individuals from distinct populations and admixed individuals nicely as also verified by a Procrustes analysis obtaining a RMSD of 0.00121, when compared to the PCA inferred from the true genotypes. There is clear bias in the results of the ngsTools model where the patterns are representing sequencing depth instead of population structure as made apparent in Figure S1. The individuals are acting as a gradient towards the origin due to their varying sequencing depth. The biased performance of ngsTools is also reflected in the corresponding Procrustes analysis with a RMSD of 0.0174.

**Figure 1.**
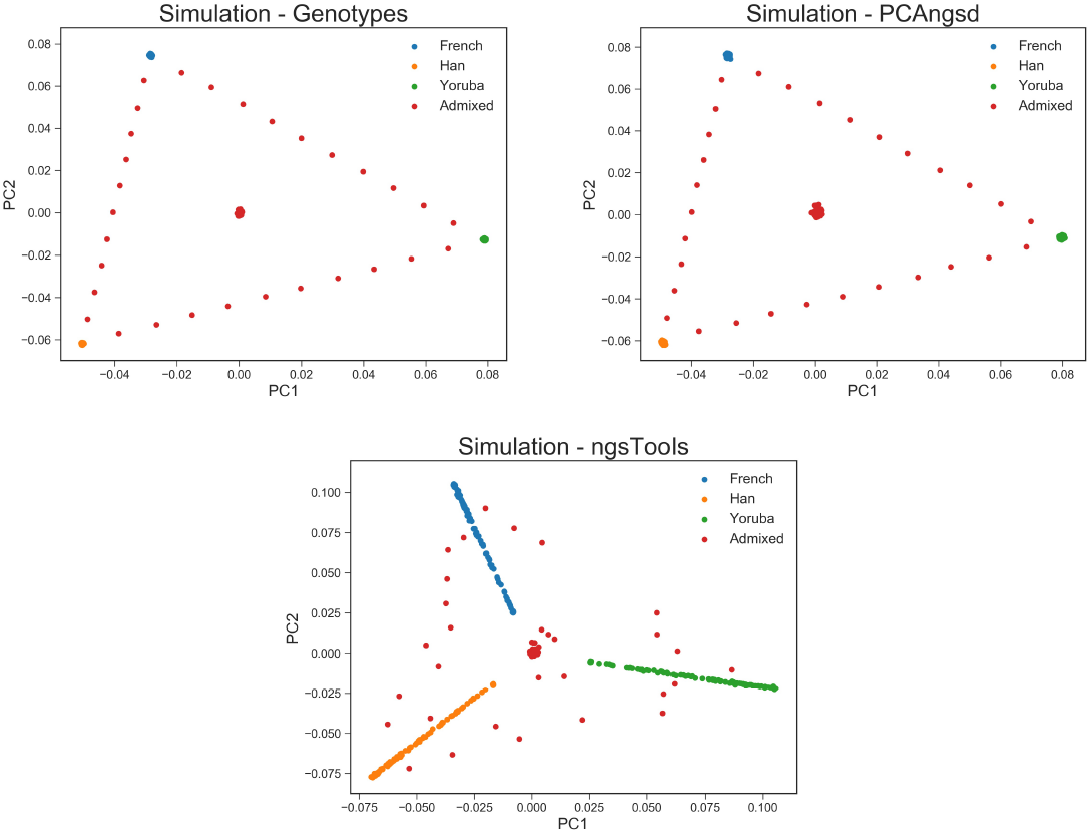
PCA plots of the top 2 principal components in the simulated dataset consisting of 380 individuals and 0.4 million variable sites. The left-hand plot shows the PCA performed on the known genotypes using equation 2. The middle plot shows the PCA performed by PCAngsd, and the righthand plot displays the PCA performed by the ngsTools model (equation 3).

The estimated admixture proportions of the simulated dataset are displayed in Figure 2. PCAngsd estimates the admixture proportions well with a RMSD of 0.00476 compared to the ADMIXTURE estimates of the known genotypes, but is however outperformed by NGSadmix with a RMSD of 0.00184. For the 380 individuals and 0.4 million SNPs using *K* = 3, PCAngsd had an average run-time of only 2.9 minutes while NGSadmix had an average run-time of 7.9 minutes.

**Figure 2.**
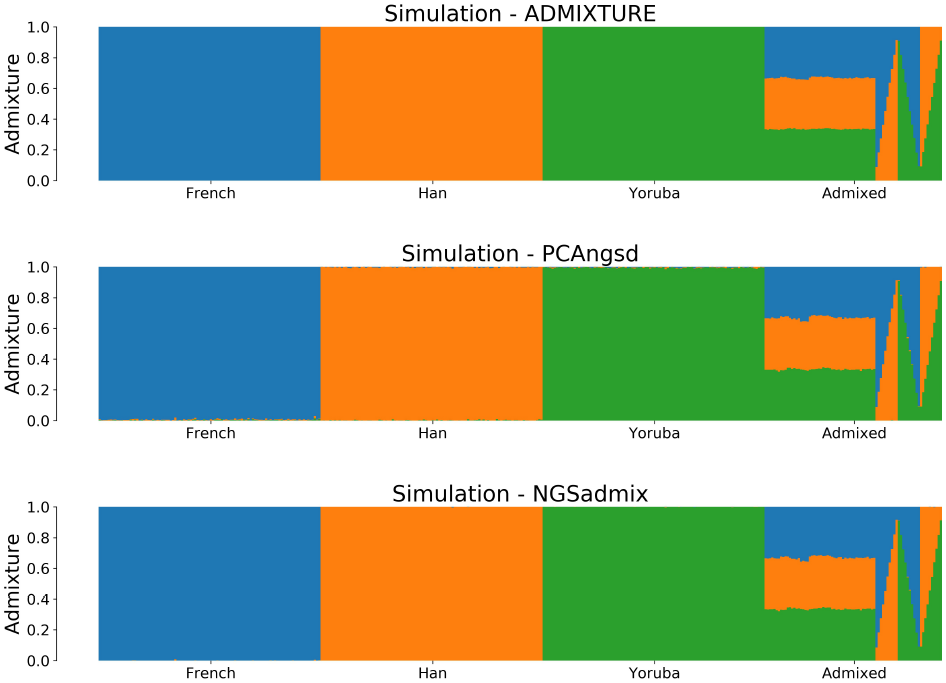
Admixture plots for *K* = 3 of the simulated dataset where each bar represents a single individual and the different colors reflect each of the *K* components. The first plot is the admixture proportions estimated in ADMIXTURE using the known geno types, which we use as the ground-truth in our simulation studies. The second plot shows admixture proportions estimated using PCAngsd with parameter *α* = 0 and the bottom plot using NGSadmix.

### 1000 Genomes

The methods of PCAngsd have also been applied to the CEU (European ancestry), CHB (Chinese ancestry), YRI (Nigerian ancestry) and MXL (Mexican ancestry) populations of the low coverage 1000 Genomes dataset. The MAP test indicated evidence of 3 significant principal components meaning that the Native American ancestry explains enough genetic variance in the dataset to represent an axis of its own. The results of the PCA are displayed in Figure 3. As was also seen for the simulated dataset, PCAngsd is able to cluster all individuals almost perfectly, while the ngsTools model is only able to capture some of the same population structure patterns with some of the populations looking admixed. Its results are still biased by the variable sequencing depth as seen as well in Figure S2. The RMSD values of the Procrustes analyses verify the observations, where PCAngsd has a RMSD of 0.00182 compared to ngsTools with a RMSD of 0.0075.

**Figure 3.**
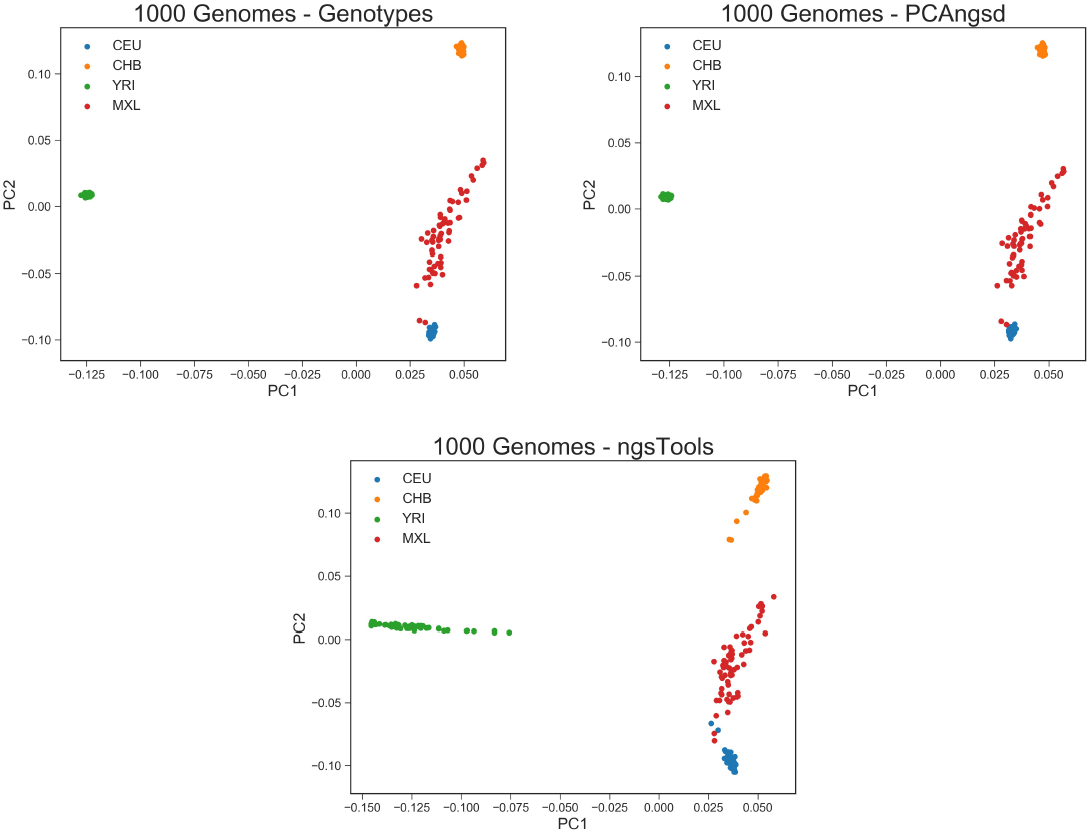
PCA plots of the top 2 principal components for the 1000 Genomes dataset with 193 individuals and 8 million variable sites. The left-hand plot is based on the known genotypes of the overlapping variable sites in the low depth NGS data, the middle plot is performed by PCAngsd and the righthand plot is performed by the ngsTools model.

The admixture plots are displayed in Figure 4. PCAngsd is not able to outperform NGSadmix in terms of accuracy, however it is still able to estimate a very similar result. PCAngsd has some issues with noise in its estimation but is however able to reduce it with the use of the sparseness parameter *α* = 1500. The likelihood measure in equation 19 has been used to easily find an optimal *α* as seen in Figure S5. PCAngsd estimates the admixture proportions with a RMSD of 0.0108 compared to NGSadmix with a RMSD of 0.007148. The average run-time for 193 individuals and 8 million SNPs using *K* = 4 was 27.3 minutes for PCAngsd and 7.1 hours for NGSadmix, making PCAngsd more than 15x faster than NGSadmix while both performing PCA and estimating admixture proportions.

**Figure 4.**
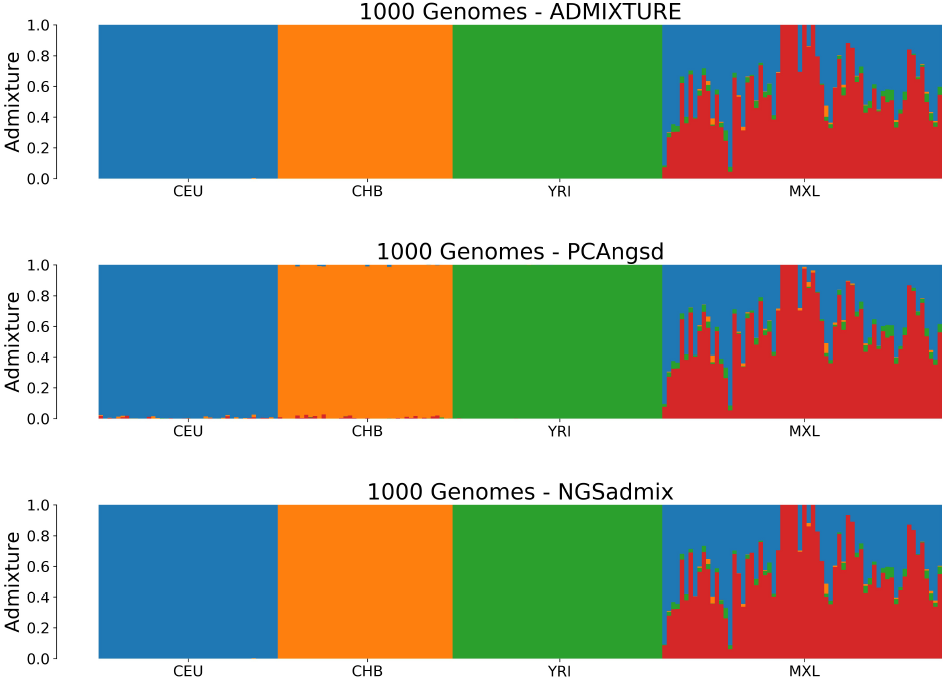
Admixture plots for *K* = 4 of the 1000 Genomes dataset where each bar represents a single individual and the different colors reflect each of the *K* components. The first plot is the admixture proportions estimated in ADMIXTURE using the known genotypes, the second plot shows admixture proportions estimated in PCAngsd with parameter *α* = 1500 and the last plot is the admixture proportions estimated in NGSadmix.

### Waterbuck

Lastly, we have analyzed the low depth whole genome sequencing waterbuck dataset consisting of 73 individuals from 5 localities. The MAP test reported 4 significant principal components for explaining the genetic variation in the dataset which also fits with having 5 distinct waterbuck sampling sites. The PCA plots are visualized in Figure 5, where the top 4 principal components have been plotted for each method. Once again, PCAngsd is able to cluster the populations much better than the ngsTools model, however the effect is not as apparent as for the other datasets. Interestingly, populations can switch positions between the two methods as seen with Samole on the second principal component and Samburu and Matetsi on the third principal component.

**Figure 5.**
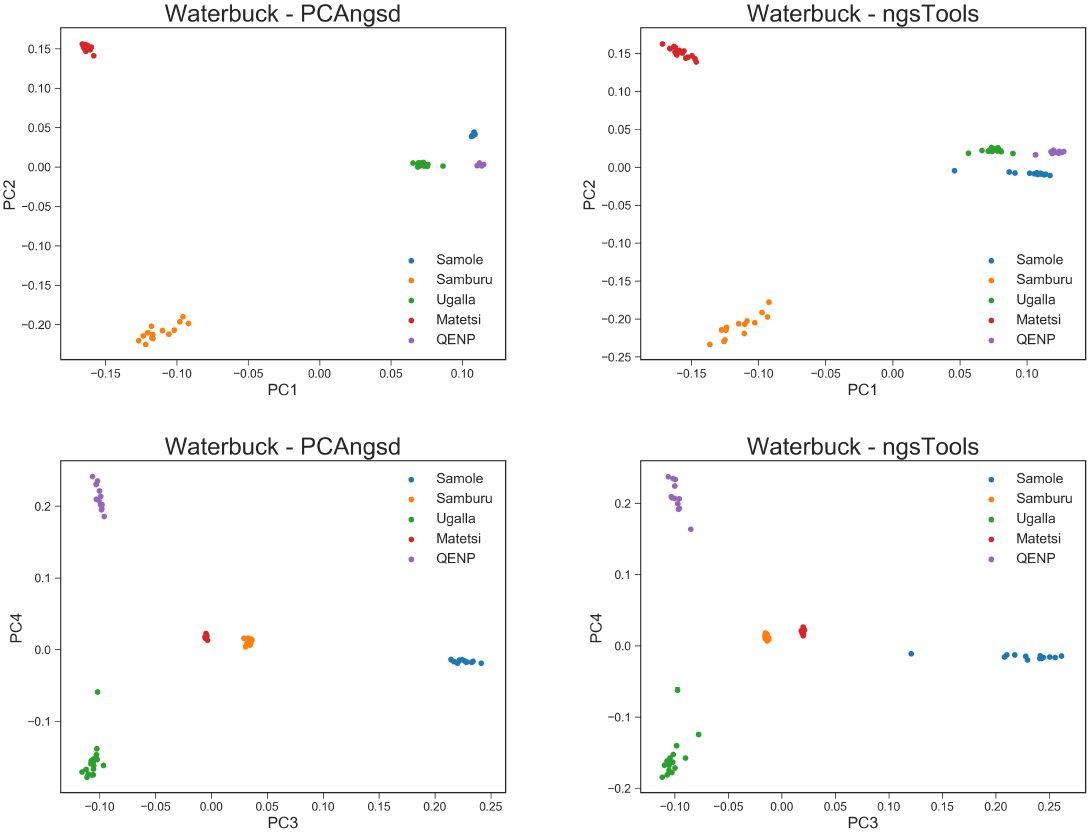
PCA plots of the top 4 principal components for the waterbuck dataset with 73 individuals and 9.4 million variable sites. The first row displays the plots of the first and second principal components for PCAngsd and the ngsTools model, respectively, while the second row displays the plots of the third and fourth principal components.

As some few clusters are not so well defined, they will affect the admixture plots seen in Figure 6, where the increased level of noise is hard to remove without also affecting the true ancestry signals. Still, PCAngsd is capturing the same ancestry signals as NGSadmix with the use of the sparseness parameter. It is worth noting that an admixed individual of Ugalla and QENP is captured in both PCA and admixture estimation of PCAngsd as also verified by the NGSadmix method. The run-times for the waterbuck dataset consisting of 73 samples and 9.4 million SNPs using *K* = 5 was an average of 14.5 minutes for PCAngsd while NGSadmix had an average run-time of 3.2 hours, thus making PCAngsd more than 13x faster.

**Figure 6.**
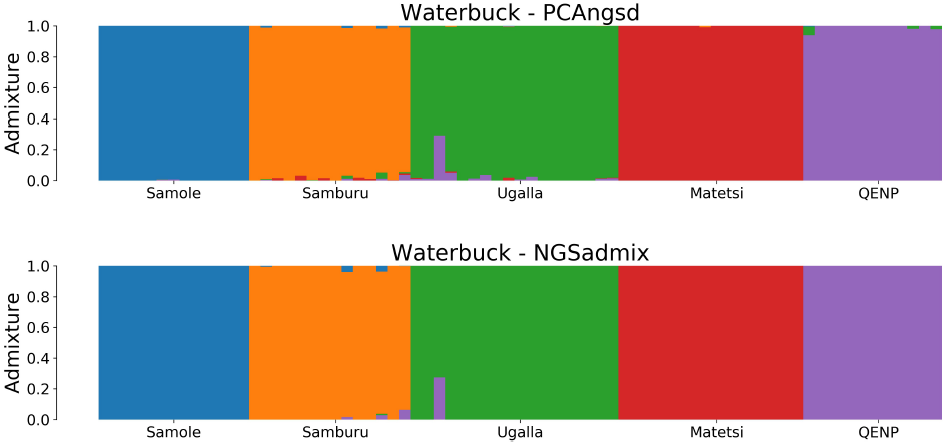
Admixture plots for *K* = 5 of the waterbuck dataset where each bar represents a single individual and the different colors reflect each of the *K* components. The first plot is the admixture proportions estimated in PCAngsd with parameter *α* = 5000 and the second plot shows the admixture proportions estimated in NGSadmix.

## Discussion

We have presented two methods for inferring population structure and admixture proportions in low depth NGS data and both methods have been implemented in a framework named PCAngsd. We have developed a framework using genotype likelihoods to iteratively estimate individual allele frequencies based on PCA. We have connected principal components to admixture proportions such that we are able to infer and estimate both in a very fast approach making it feasible to analyze large datasets.

Based on the results when inferring population structure using PCA, it is clear that the increased uncertainty of low depth sequencing data biases the clustering of populations using the ngsTools model which also takes genotype uncertainty into account. Contrary to PCAngsd, population structure is not taking into account when using the posterior genotype probabilities to estimate the covariance matrix. The ngsTools model uses population allele frequencies as prior information for all individuals such that individuals are assumed to be sampled from a homogeneous population. This assumption is of course violated when individuals are sampled from structured populations with diverge ancestries. Missing data is therefore modeled by population allele frequencies that resemble an average across the entire sample, which is similar to setting standardized genotypes to 0 in the estimation of the covariance matrix for genotype data. As an effect of this, the low depth individuals are modeled by sequencing depth instead of population structure. These results may lead to misinterpretations of population structure or admixture only due to low and variable sequencing depth. But the bias is not seen for individuals with equal sequencing depth as shown in Figure S4 for the ngsTools model. Here all individuals have been simulated with an average sequencing depth of 2.5 𝒳 such that individuals will approximately inherit the same amount of missing data. However, PCAngsd is able to overcome the observed bias of low and variable sequencing depth by using individual allele frequencies as prior information, which leads to more accurate results in all datasets of the study, as missing data is modeled by inferred population structure. The assumption of conditional independence between individuals in the estimation of the covariance matrix (equation 10) also holds for structured populations by conditioning on individual allele frequencies.

The number of significant eigenvectors used in the estimation of individual allele frequencies is determined by the MAP test. The MAP test is performed on the covariance matrix estimated from the ngsTools model. Thus in cases of complex population structure and low and variable sequencing depth, it is possible that the MAP test will not find a suitable number of significant eigenvectors to represent the genetic variation of the dataset. It could therefore be more relevant to use prior information regarding the number of eigenvectors needed for the dataset instead. However for each of the cases analyzed in this study, the MAP test inferred the expected number of significant eigenvectors to describe the population structure.

PCAngsd is able to approximate the results of NGSadmix to a high degree when estimating admixture proportions using solely the estimated individual allele frequencies. However, PCAngsd is not able to outperform NGSadmix in terms of accuracy, but it is however able to capture the exact same ancestry patterns as the clustering-based methods in a much faster approach, as shown by the run-times of each method. Another advantage of PCAngsd is that the estimated individual allele frequencies are only needed to be computed once for a specific *K*, thus multiple different *α*’s and random seeds can be tested in the same run for an even greater speed advantage over NGSadmix, since the iterative estimation of individual allele frequencies is the most computational expensive step in PCAngsd. PCAngsd is therefore an appealing alternative for estimating admixture proportions for low depth NGS data as convergence and run time can be a problem for a large number of parameters in NGSadmix. PCAngsd was only seen to converge to a single solution for all our practical tests, where we used five batches for all analyses (*B* = 5).

**Table 1.** Average run-times of 10 initializations for both PCAngsd and NGSadmix. The run-times reported for PCAngsd include reading of data and estimation of covariance matrix and admixture proportions, while run-times listed in parentheses only include estimation of admixture proportions, when parsing previously estimated individual allele frequencies. All tests have been performed server-side using 32 threads.

| Dataset | n | m | K | PCAngsd | NGSadmix | Depth |
| --- | --- | --- | --- | --- | --- | --- |
| Simulated | 380 | 0.4 million | 3 | 2.9 min (2.1 min) | 7.9 min | 0.5 – 5X |
| 1000 Genomes | 193 | 8 million | 4 | 27.3 min (19.5 min) | 424.9 min | 1.5 – 12.5X |
| Waterbuck | 73 | 9.4 million | 5 | 14.5 min (9.3 min) | 192 min | 2.2 – 4.7X |

Both methods of the PCAngsd framework rely on an representative set of individual allele frequencies which we model using the inferred principal components of the SVD on the genotype dosages. The number of individuals representing each population or subpopulation is essential for inferring principal components that describe true population structure as each individual will contribute to the construction of these axes of genetic variation. This particular effect can be seen in the PCA results of the waterbuck dataset where the populations are only described by a low number of individuals such that some of the clusters are not so well defined as for the other datasets. The admixture proportions estimated from the waterbuck dataset are therefore affected as well which can be seen by the additional noise in the admixture plots.

The PCAngsd framework might be able to push the lower boundaries of sequencing depth required to perform population genetic analyses using NGS data of largescale genetic studies. PCAngsd demonstrates an effective approach on dealing with merged datasets of various sequencing depths as well, as missing data will be modeled by population structure. The estimated individual allele frequencies contain a lot of information regarding population structure and can open up for the development and extension of population genetic models based on a similar probabilistic framework to naturally correct for population structure in order to obtain more accurate estimates in heterogeneous populations.

